# Sequential invasion of epithelial and immune cells in Shigellosis

**DOI:** 10.64898/2026.09.11.751053

**Authors:** Niti B. Jadeja, Brittany A. DeVasure, Lauren K. Yum, Hervé F Agaisse

## Abstract

*Shigella flexneri*, the agent of shigellosis, invades the colonic mucosa, which leads to mucosal erosion, blood and immune cell infiltration, culminating in the hallmark of the disease, bloody diarrhea. Here, we conducted a microscopy-based analysis of time-resolved infection experiments in the infant rabbit model of shigellosis. We observed two stages of infection in the colonic tissue. During the early stage of infection, bacteria invaded and spread in epithelial cells, which correlated with mucosal erosion and blood infiltration. During the late stage of infection, the bacteria transitioned from their primary niche, the epithelial compartment, to their secondary niche, the lamina propria. Transcriptomic analysis of the late stage of infection assigned identities to various epithelial, immune, and stromal cells. The vast majority of the bacteria were associated with neutrophils or macrophages during the late stage of infection. We provide a discussion of how this model compares and contrasts with previous models of shigellosis.

## Introduction

*Shigella* spp., mainly *S. flexneri* and *S. sonnei*, continue to impose a substantial global burden, causing over 250 million cases of bacillary dysentery annually (bloody diarrhea due to *Shigella* spp. infection), resulting in approximately 200,000 deaths, including 63,713 deaths in children younger than 5 years^1^. In the United States, 450,000 infections are reported annually, which account for more than $90 million in direct medical costs^2^. The lack of a licensed vaccine and the increasing threat of antimicrobial resistance in *Shigella* spp. underscore the need to address the remaining knowledge gaps in our understanding of the mechanisms supporting the disease^3^.

Seminal studies conducted in non-human primates have revealed that *S. flexneri* is an intracellular pathogen that resides in epithelial cells in the colon^4^. The development of *in vitro* tissue culture systems^5^ was instrumental in understanding the molecular determinants supporting *S. flexneri* intracellular infection. *S. flexneri* invasion of epithelial cells relies on the presence of “the invasion plasmid”^6^ that harbors the 37kb “entry region”^7^ encoding the type-3 secretion system (T3SS). The T3SS effector proteins manipulate various cellular processes, including the actin cytoskeleton, leading to the uptake of the bacteria by non-phagocytic cells, such as epithelial cells^8^. *S. flexneri* quickly escapes primary vacuoles and gains access to the cytosol of infected cells. Cytosolic bacteria display actin-based motility and spread from cell-to-cell^9,10^. The dissemination process is critical for *S. flexneri* pathogenesis, as spreading defective mutants are basically avirulent in non-human primates^11^ and infant rabbits^12^.

The current model of shigellosis that reached the textbooks was established in an effort to provide an integrated view based on observations made in tissue culture systems constituted of human polarized epithelial cells or murine macrophages, and an animal model using the small intestine of adult rabbits. A first key observation has been the fact that *S. flexneri* cannot invade polarized cells through their apical side *in vitro*^13^. However, treatments that relaxed lateral junctions led to massive infection. This led to the concept that *S. flexneri* must invade epithelial cells through their basal and/or lateral sides^13^. The second key observation came with early studies using the ligated ileal loop of adult rabbits, showing that the primary sites of infection are M cells in the Peyer patches^14^. Transcytosis through M cells delivers *S. flexneri* directly into the Lamina Propria, where bacteria encounter immune cells. *S. flexneri* interaction with macrophages leads to cell death *in vitro* and *in vivo*^15,16^. Dead macrophages release bacteria at the basal pole of the epithelium, thereby promoting epithelial invasion and subsequent cell-to-cell spread. These events allow bacteria to colonize their main niche: the colonic epithelium. Macrophage death also results in the release of inflammatory molecules^17,18^ leading to the massive recruitment of neutrophils that clear the infection, but also chiefly contribute to tissue destruction^19,20^. Altogether, these observations promoted the widespread notion that shigellosis is a neutrophil-driven inflammatory disease^21^.

Although the intestinal loop model was instrumental in understanding various aspects of the inflammatory response occurring upon infection, the small intestine is not the site of infection in humans, and animals do not display bloody diarrhea in the intestinal loop model. In an effort to address these important caveats, we have developed a new model of shigellosis in the colon of infant rabbits^12^. Importantly, this model does recapitulate all symptoms of human shigellosis, including severe epithelial fenestration, vascular lesions, and immune cell infiltration, leading to profuse bloody diarrhea. As demonstrated below, the new model offers a different perspective on the sequence of events occurring during colonic infection.

## Results

### Time course analysis of bacterial colonization

To characterize the spatial and temporal invasion of the colonic mucosa during acute shigellosis in infant rabbits (0-24 hours post-inoculation, hpi), we determined the timeline of bacterial burden using fluorescence microscopy. To this end, we harvested distal colons of infected animals at 4, 8, and 24 hpi along with PBS-treated uninfected controls. We conducted immuno-quantification and image analysis of the colonic tissue stained for *S. flexneri* and counter-stained for the epithelial marker E-cadherin (Supplementary Fig. 1), which revealed the evolution of bacterial burden across the experimental timeline (Fig. 1). During the early stages of infection (4 and 8 hpi), animals challenged with *S. flexneri* 2457T displayed a clear presence of bacteria within the mucosa (Fig. 1. a-c). By 24 hpi, bacterial burden had significantly increased but stayed confined to the mucosa, never crossing into the sub-mucosa (Fig. 1e). Interestingly, *S. flexneri* was strongly associated with E-cadherin-positive cells (ECs) during the early phase of infection (Fig. 1d,f, EC). By 24 hpi, however, *S. flexneri* was no longer associated with E-cadherin-positive ECs (Fig. 1e) and was mainly associated with the lamina propria (LM) (Fig. 1f, LM). High-resolution imaging determined that the vast majority of bacteria associated with the lamina propria existed in non-E-cadherin-expressing cells (NECs) during late infection (Supplementary Fig. 2). These data indicate that, in the infant rabbit colon, infection begins with invasion of ECs, propagates through cell-to-cell spread within the epithelial niche during the early stage of infection, and then disseminates during the late stage of infection to the lamina propria where bacteria are associated with NECs, presumably immune cells.

**Fig. 1.**
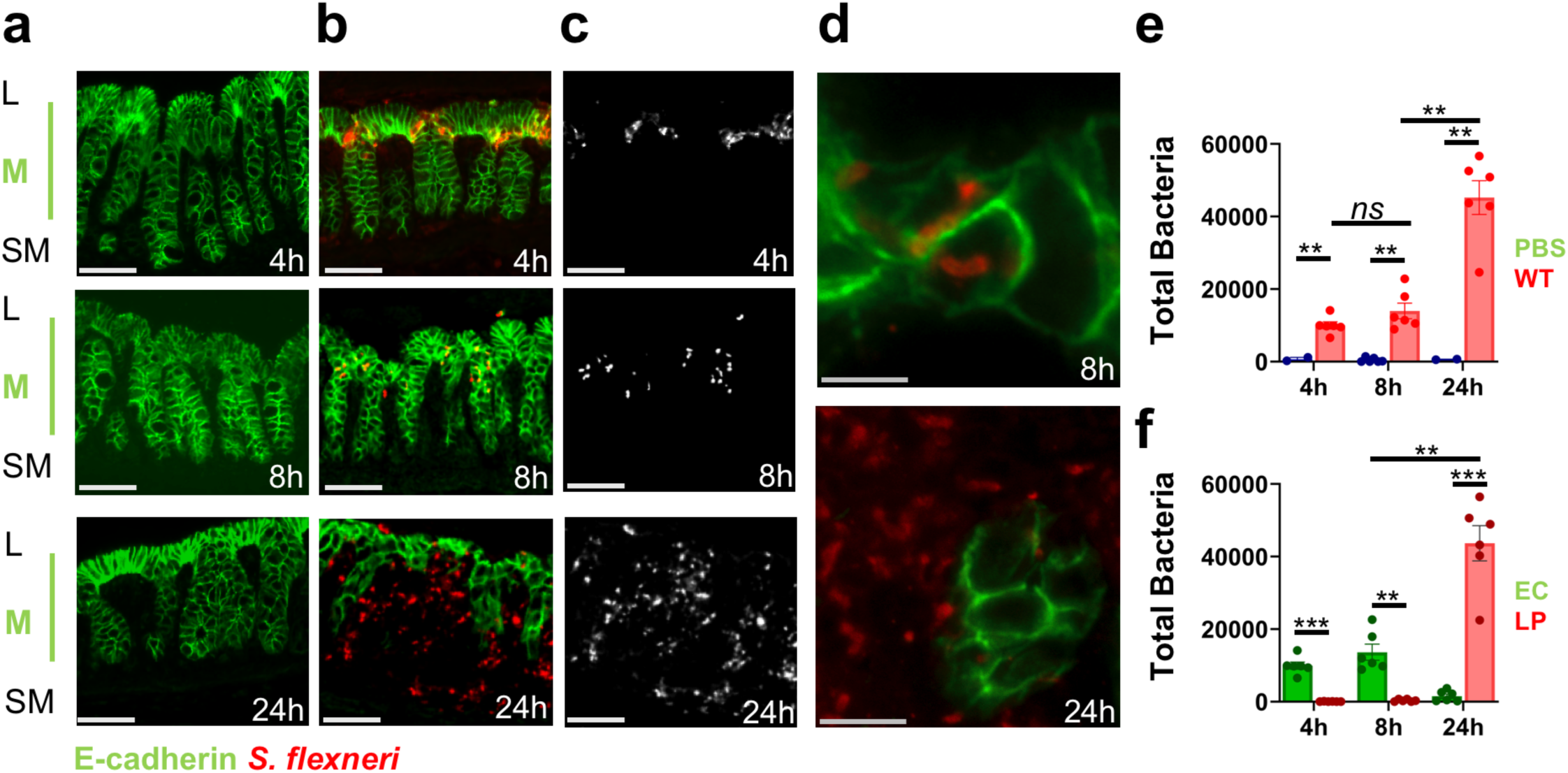
Time course of *S. flexneri* infection in the infant rabbit colon. (**a,b**) Representative immunofluorescence images of colonic sections from animals treated with PBS (a) or *S. flexneri* (b) at the indicated time points, stained for E-cadherin (green) and *S. flexneri* (red). L, lumen; M, mucosa; SM, submucosa. **(c)** Bacterial channel from the merged images as shown in (b). **(d)** High-magnification images of infected tissue at 8 and 24 hpi, showing bacterial localization relative to E-cadherin-positive cells. Scale bars, 100 µm (a-c) and 10 µm (d). **(e)** Bacterial burden per colon in mock-treated and infected tissue. Counts represent the sum of images acquired per colon. Bars show mean ± SEM with individual biological replicates overlaid. Infected animals, n=6 colons per time point; PBS treatment, n=3 (4 hpi) and n= 4 (8 and 24 hpi). Total intracellular bacterial burden was clearly detected in WT-infected tissue over background in PBS-treated controls 4 hpi (9,974 ± 987 vs. 24 ± 12 bacteria/colon, *p*= 0.0012), 8 hpi (13,978 ± 2,167 vs. 30 ± 17, *p*= 0.0091), and 24 hpi (45,221 ± 4,648 vs. 44 ± 18, *p*= 0.0014). In infected animals, counts increased significantly between 4 and 24 hpi and 8 and 24 hpi (*p*= 0.0035 and *p*= 0.0040), with no significant change between 4 and 8 hpi (*p*= 0.58). Because within-group variance scaled with the mean, groups were compared by Brown-Forsythe and Welch ANOVA with the Games-Howell post-test (15 pairwise comparisons, α = 0.05). A two-way ANOVA on the same data confirmed a significant time × treatment interaction (F (2, 23) = 24.08, *p*< 0.0001). \*\**p*< 0.01; ns, not significant). **(f)** Total bacterial burden partitioned into E-cadherin-positive epithelial cells (EC) and E-cadherin-negative Lamina Propria (LP). Bar graphs show total intracellular bacteria in epithelial (EC) and lamina propria (LP) compartments at 4, 8, and 24 hpi. At 4 and 8 hpi, bacterial burden was significantly higher in EC than LP (4 hpi: 9,934 ± 994 vs. 40 ± 21 bacteria/colon, *p*= 0.0005; 8 hpi: 13,647 ± 2,221 vs. 331 ± 137, *p*= 0.0021). By 24 hpi, bacterial burden was greater in LP than in EC (43,672 ± 4,832 vs. 1,549 ± 595, *p*= 0.0008). LP counts increased significantly between 8 and 24 hpi (*p*= 0.002), with no significant change between 4 and 8 hpi (*p*= 0.41). Data represent mean ± SEM. EC and LP were compared within each time point by paired t-test with Holm-Šídák correction; comparisons across time points were made by Brown-Forsythe and Welch ANOVA with the Games-Howell post-test. \**p*< 0.05; \*\**p*< 0.01; \*\*\**p*< 0.001; ns, not significant.

### Early events during the colonization process

The current model of shigellosis stipulates that *S. flexneri* invade M cells first, and quickly transcytose to the lamina propria compartment where they infect and kill macrophages in less than 4 hours^16^. To determine early events of infection in the infant rabbit model, we infected animals for 2 hours and assayed for the location of bacteria in the tissue by immunostaining. As previously shown by our group^12^, bacteria gained access to the epithelial compartment within the two-hour window of the infection process (Fig. 2a,b). Moreover, very few bacteria gained access to the lamina propria (Fig. 2a,b). As previously described for other models of shigellosis^16^, we used the TUNEL assay to quantify cell death in the mucosa. The early time points displayed no significant cell death relative to PBS treatment control up to 8 hpi (Fig. 2c,d). The only time point tested that displayed significant cell death was the 24-hour time point, when bacteria gained access to the lamina propria and potentially interacted with macrophages (Fig. 2c,d).

**Fig. 2.**
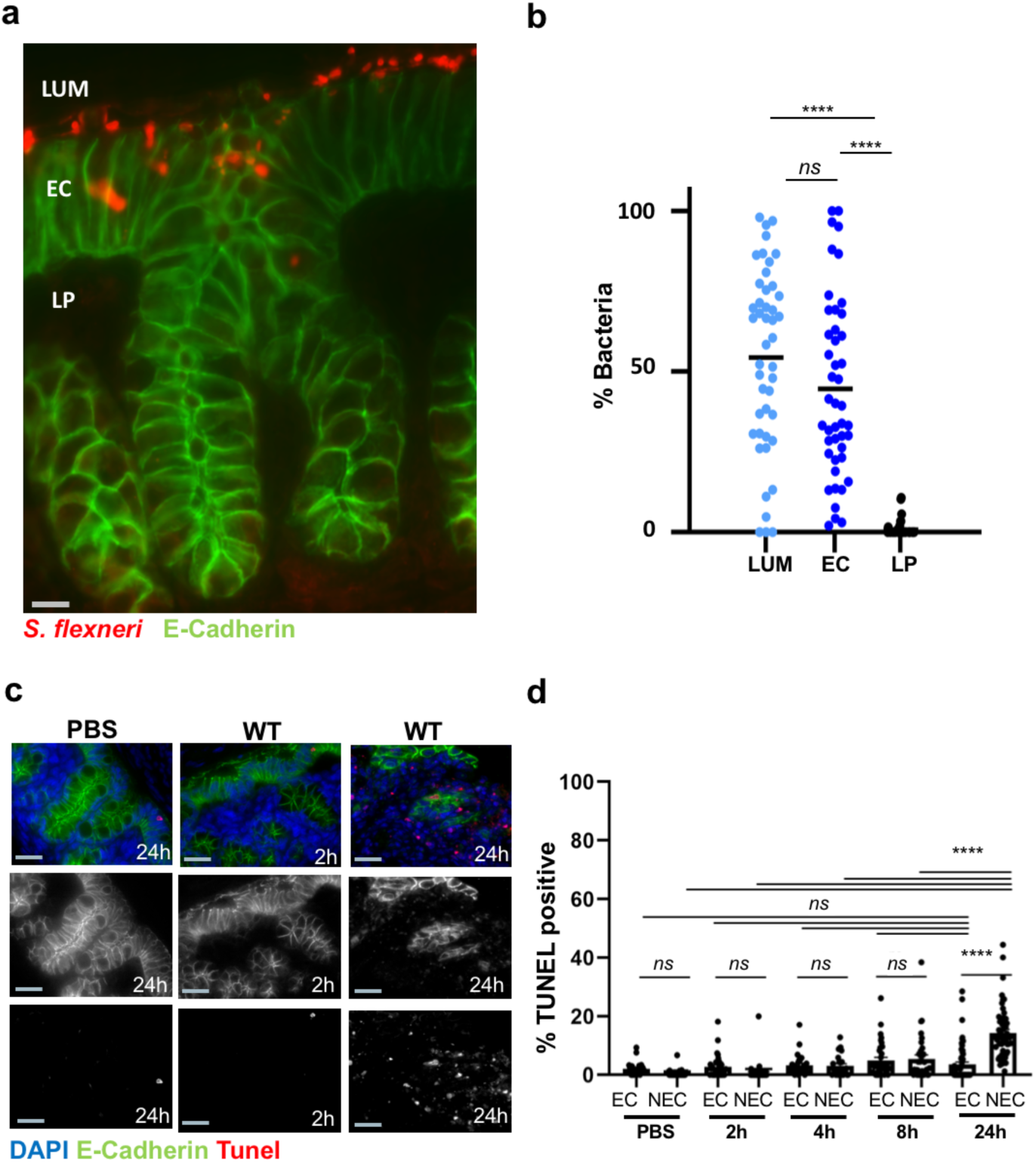
Early stage of bacterial invasion on the colonic mucosa. **(a)** Representative image of colonic section 2 hpi, immuno-stained for E-cadherin (green) and *S. flexneri* (red), showing 3 compartments: outside the tissue (LUM), intracellular (EP) and lamina propria (LP). Scale bar: 10 µm. **(b)** Quantification of bacteria localized to each compartment in E-cadherin-positive cells. Bars show group means with individual images overlaid. Bacteria were largely confined to the outside and the intracellular EC compartment, with few detected in the LP. Bacterial counts were significantly higher in EC than LP (47.4 ± 5.2 vs. 1.8± 0.5) whereas LUM and EC counts did not differ significantly (*p*= 0.18). Because group variances differed significantly (F (19,19) = 26.15, *p*< 0.0001), groups were compared by unpaired two-tailed *t*-test with Welch’s correction (t = 17.86, df = 20.45, *p*< 0.0001). \*\*\*\**p*< 0.0001; ns, not significant. **(c)** Representative images of TUNEL staining on sections of infant rabbit colons treated with PBS or *S. flexneri* at 2 and 24 hours. Scale bars, 50 µm. **(d)** TUNEL staining revealed EC cell death did not differ significantly between any two time points (all pairwise comparisons *p*> 0.05; Welch’s *t*-test with Holm-Šídák correction across six comparisons). In contrast, LP cell death was indistinguishable from PBS at 2 hpi (0.9 ± 0.6%, P = 0.76), became significantly elevated at 4 and 8 hpi (3.1 ± 0.7%, P = 0.0005; 5.5 ± 1.3%, P < 0.0001), and increased markedly by 24 hpi (14.3 ± 1.2%, P < 0.0001 vs. PBS). EC and NEC cell death did not differ significantly at 4 or 8 hpi (both p> 0.99), although EC modestly exceeded LP at 2 hpi (p= 0.0058). By 24 hpi, NEC cell death was significantly greater than EC (14.3 ± 1.2 % vs. 3.5 ± 0.9%, p< 0.0001), representing a ∼3.5-fold difference between compartments. Data are presented as mean ± SEM. EC cell death was compared across time points by Welch’s t-test; EC and NEC were compared within each field by Wilcoxon signed-rank test; and each compartment was compared with PBS-treated controls by Mann-Whitney U test. All families were corrected by the Holm-Šídák method. *p< 0.05; **p< 0.01; ***p< 0.001; ****p< 0.0001; ns, not significant.

We also evaluated ulceration and bleeding symptoms by quantifying epithelial fenestration and red blood cell pockets in the colonic tissue on H&E staining images (Fig. 3a-d). Animals inoculated with PBS exhibited low levels of epithelial fenestration, ranging from 5-15% (Fig. 3e), potentially due to basal remodeling of the epithelial structure or damage inflicted on the tissue upon dissection. We observed no statistical differences in fenestration between PBS control and animal samples collected at 4 hpi (Fig. 3e). 8 hpi, a significantly higher level of fenestration was observed compared to PBS control and 4 hpi (Fig. 3e). By 24 hpi, the colonic tissue exhibited severe epithelial fenestration (Fig. 3e). Red blood cell infiltration started as early as 4 hpi and increased gradually up to 24 hpi (Fig. 3f).

**Fig. 3.**
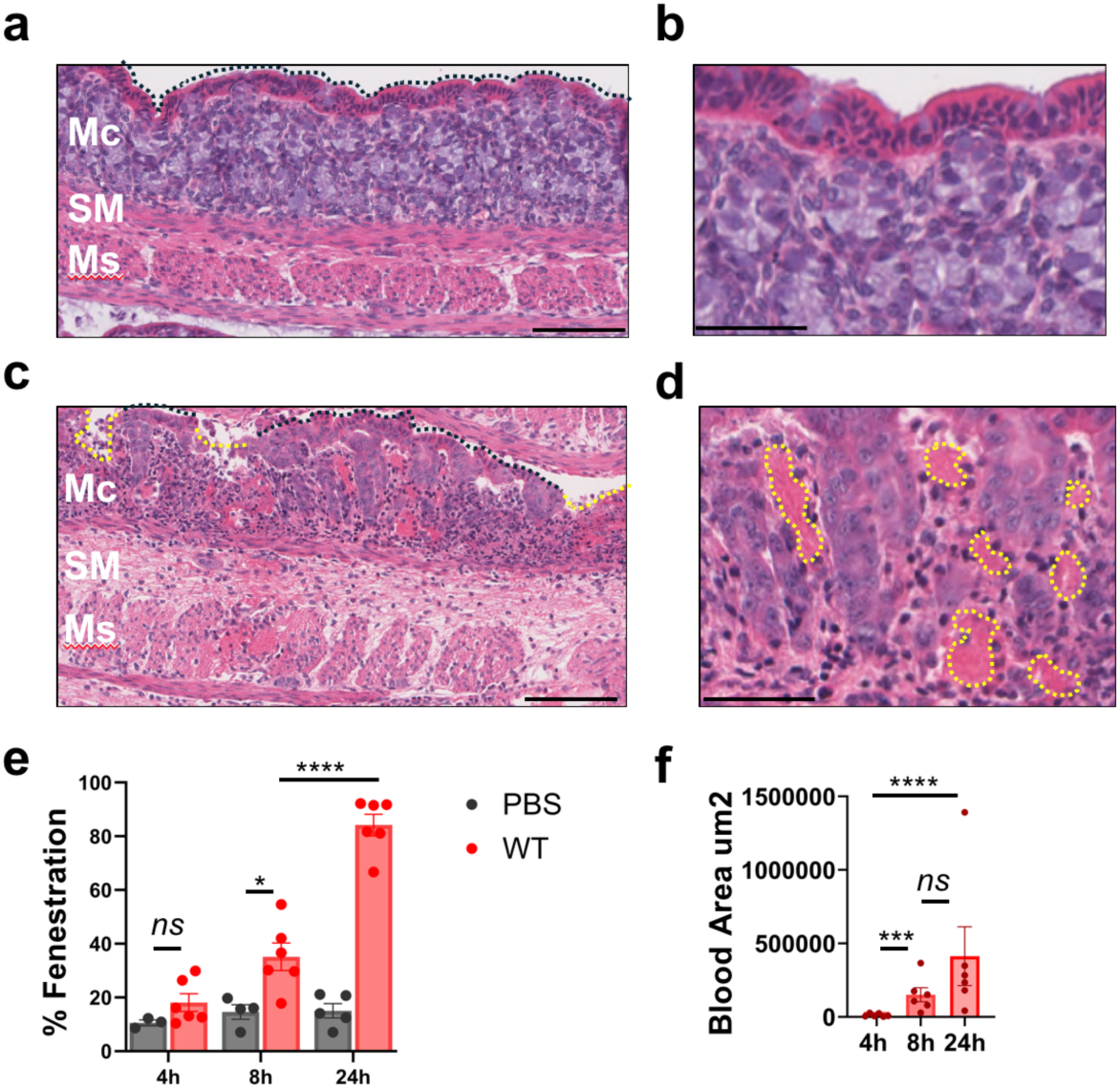
Epithelial fenestration. **(a-d)** Representative images of *S. flexneri* infection in infant rabbit colons using Hematoxylin and Eosin staining of colonic sections of animals inoculated with PBS **(a, b)** or **(c,d)** *S. flexneri,* depicting fenestration and blood area measured across mock and infected samples at 24hpi. Mc denotes mucosa, SM: sub-mucosa, and Ms: Muscle. Scale bar: b-d: 100 µm and c-d: 50 µm. **(e)** Percent fenestration across the infected colon length across the time course in mock and 24 hpi **(f)** Within the 24hpi infections, measure of the area of blood across the total length of the infected colon. Data represent 6 individual biological replicates with mean ± SEM. Statistical comparisons were performed using two-way ANOVA and unpaired t-tests. \**p*< 0.05; \*\*\**p*< 0.001; \*\*\*\**p*< 0.0001; ns, not significant. Source data are provided as a Source Data file available upon request.

Altogether, these experiments indicate that bacterial infection initiates in epithelial cells where bacteria grow and spread from cell to cell. This early colonization event correlates with the beginning of quantifiable damage inflicted to the tissue, including epithelial fenestration and blood cell infiltration within 8 hpi. By 24 hpi, bacteria have multiplied considerably and re-localized to the lamina propria, while the colonic tissue experienced severe epithelial fenestration and blood infiltration leading to bloody diarrhea.

### Transcriptomic analysis of host responses in the infected colon

To further characterize the host response to shigellosis, we analyzed the host transcriptional responses to infection 24 hpi by conducting RNA-sequencing experiments. To this end, we isolated total RNA from the colon of infant rabbits 24 hours after PBS treatment or infection with *S. flexneri* 2457T. We next conducted bulk RNA-Sequencing analysis using Illumina platforms and mapped the generated outputs to the available rabbit reference genome (mOryCun1.1, GCF_964237555.1). Global gene expression was determined for each sample and comparison conducted using the differential gene expression package DESeq2. The principal component analysis revealed that the transcriptional profiles of control and wild-type samples clustered separately, highlighting differences representing infection-mediated changes either in transcription levels of a given cell population or changes in the cell population composition in the colonic tissue (Fig. 4a).

**Fig. 4.**
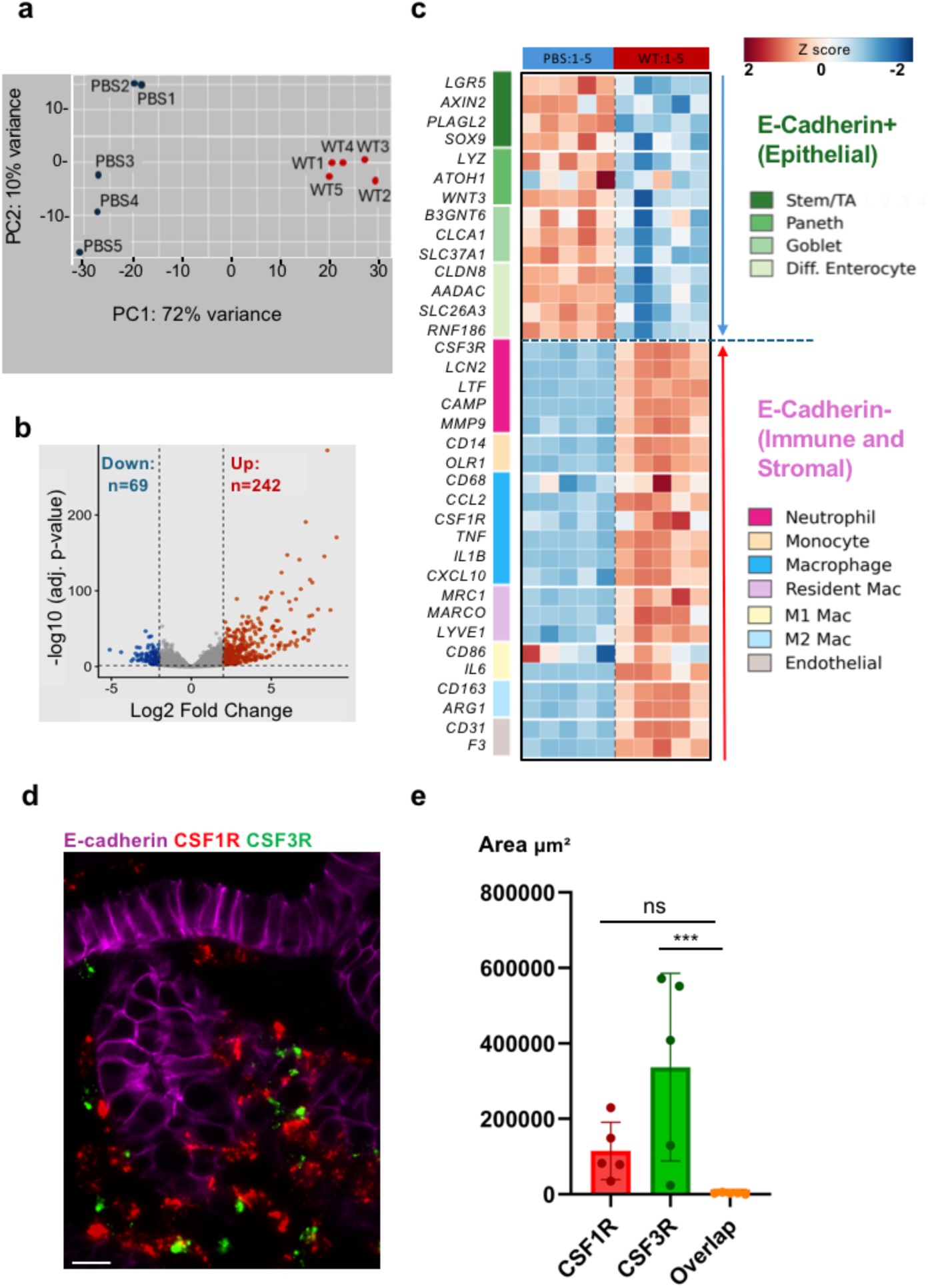
Transcriptomic analysis of the late phase of shigellosis. (**a**) Principal component analysis of rlog-transformed expression values for colonic tissue from PBS-treated (n = 5) and *S. flexneri*-infected (24 hpi; n= 5) animals. (**b**) Volcano plot of differentially expressed genes at 24 hpi vs PBS. Dashed lines indicate the thresholds applied (adjusted P < 0.05, log2 fold change >2) (**c**) Heat map of rlog-transformed read counts for differentially expressed genes that correspond to cell-type signatures defined in the MSigDB C8 cell type signature collection. Genes are grouped as E-cadherin-positive (epithelial) or E-cadherin-negative (immune/stromal) populations. Red denotes higher and blue lower relative expression (Z score). **(d)** Representative high-magnification images of signals corresponding to the neutrophil– and macrophage-specific markers CSF3R (green) and CSF1R (red), respectively, in the colonic mucosa counter-stained for E-cadherin (purple). Scale bar: 50 µm. **(e)** Quantification of CSF1R+, CSF3R+, and CSF1R+/CSF3R+ immune cells as shown in **(a)**. Bars show mean ± SD; each dot represents one colon (n=4).

Accordingly, in reference to PBS treatment, infected samples exhibited a dramatic increase in transcriptomic responses as assessed by values above or below 2-fold change (Fig. 4b, volcano plot). The normalized gene counts from the differential gene analysis were used with the C8 database derived from single-cell RNA-seq studies, defined into gene sets for specific human cell types identified in tissues. We undertook deconvolutions for our bulk RNA-Seq, analyzing cell type markers enriched at 24 hpi. As expected from the observed decrease in ECs, genes displaying significant decrease in abundance corresponded to epithelial cell types including stem and transit amplifying cells, Paneth cells, goblet cells and differentiated colonocytes (Fig. 4c). Genes increased in abundance, whose functions largely mapped to inflammatory processes reported on the infiltration/activation of various immune cells including neutrophils, monocytes, macrophages, resident macrophages, M1 macrophages, M2 macrophages, as well as endothelial cells (Fig. 4c). We experimentally profiled the immune cell infiltration by FISH (RNA-scope) with specific probes targeting the messenger RNA of CSF3R and CSF1R, two pan markers of human neutrophils and macrophages, respectively^22,23^ (Figure 4d). The two markers were highly expressed and, as expected, did not overlap (Fig. 4e).

### Identification of macrophages and neutrophils in the colonic mucosa

We next used *cell detection* modules in QuPath^24^ to determine the boundaries of NECs in the lamina propria using the DAPI nuclear stain as a reference (Fig. 5a-d NEC mask). Neutrophil– and macrophage-specific FISH probes for CSF3R and CSF1R expression were used in combination with *S. flexneri* immuno-staining. The approach showed that (25 ± 3)% and (65 ± 3)% of NECs expressed the CSF1R and CSF3R markers, respectively (Fig. 5e,f). We also tested the expression of known neutrophil– and macrophage-specific markers supporting the response to infection in humans, such as the neutrophil gelatinase-associated lipocalin LCN2^25^ and the scavenger receptor MARCO^26,27^. The approach showed that (10 ± 2)% of CSF1R-positive NECs expressed the macrophage-specific marker MARCO, whereas (40 ± 3)% of CSF3R-positive NECs expressed the neutrophil-specific marker LCN2 (Fig. 5e,f). In agreement with the notion of specificity, (95 ± 2)% of MARCO-positive and (99 ± 0.3)% of LCN2-positive NECs were macrophages and neutrophils, respectively. These results indicate that the vast majority of NECs present in the mucosa 24 hpi are CSF3R-positive neutrophils and CSF1R-positive macrophages that, similar to their human counterpart express specific markers, such as LCN2^25^ and MARCO^26,27^, respectively.

**Fig. 5.**
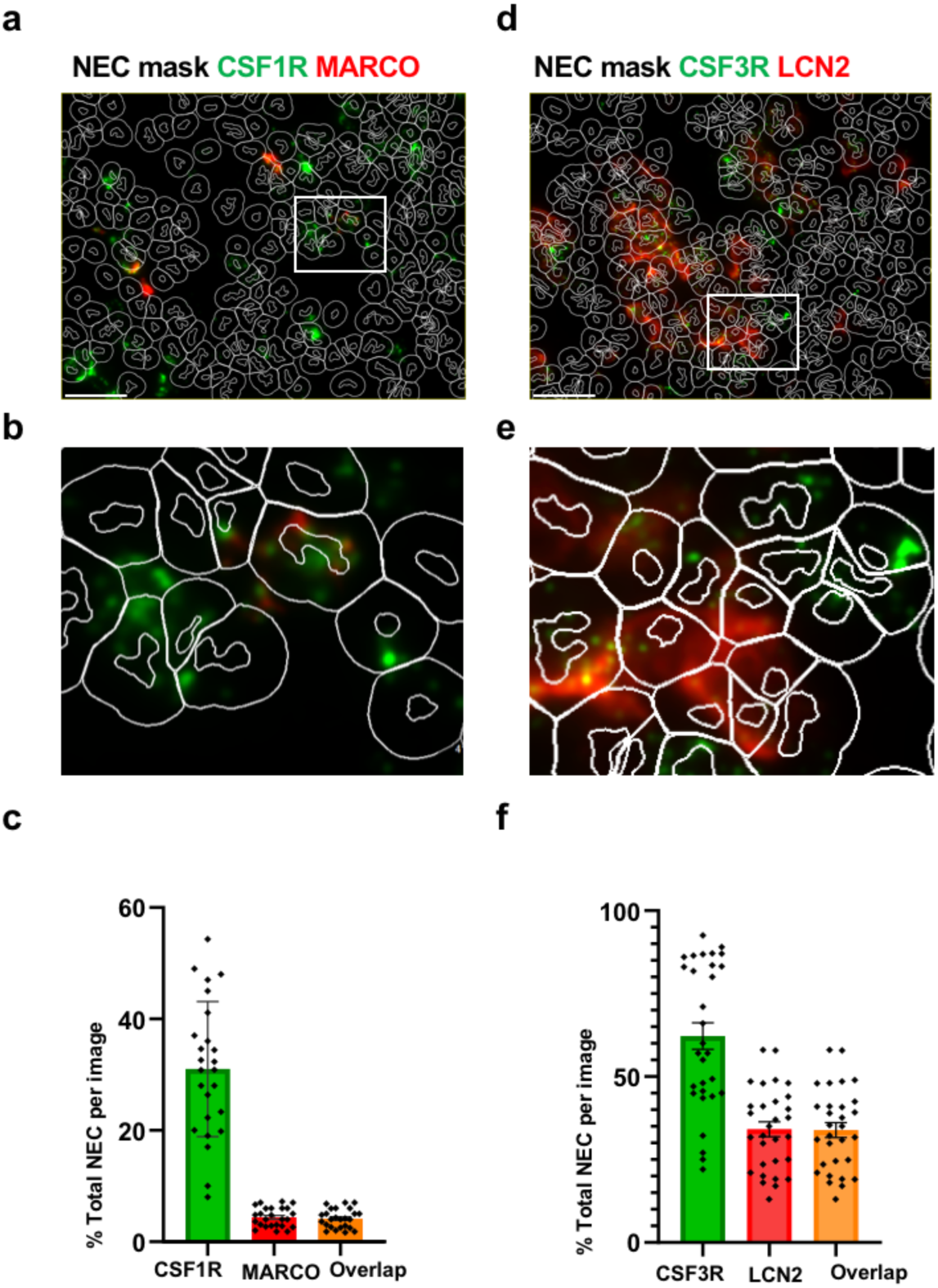
Identification of macrophages and neutrophils in the colonic mucosa. Representative images of colonic mucosa 24 hpi showing NECs (Mask, white) stained for CSF1R (a-b, green) and CSF3R (d-e, green) co-stained with a secondary marker MARCO (a-b, red) and LCN2 (d-e, red). Scale bar: 100 µm (c) Quantification of NECs that are CSF1R+ cells, MARCO+ cells and CSF1R+MARCO+ cells in colonic tissue. (f) Quantification of CSF3R+ cells, LCN2+ cells and CSF3R+LCN2+ cells in colonic tissue. Each symbol represents one image; bars show the mean and error bars, SEM (n= 25-30 images). All measures are expressed as a percentage of DAPI+ nuclei in the same image. No statistical comparison is shown between bars, as the co-positive population is a subset of each single-marker population; co-positivity instead exceeded that expected under independent expression by 3 fold (observed 4.0% vs expected 1.4%, *p*< 0.001 in (c) and by 1.5-fold (observed 33.8% vs expected 23.3%, *p*< 0.001) in (f).

### *S. flexneri* colonizes neutrophils and macrophages during the late phase of infection

We next characterized the infected NECs in the lamina propria during the late phase of infection (Supplementary Fig. 2). Neutrophil– and macrophage-specific FISH probes for CSF3R and CSF1R expression were used in combination with *S. flexneri* immuno-staining (Fig. 6). High-resolution imaging of colonic sections and computer-assisted image analysis established that (89 ± 1)% of NECs were associated with *S. flexneri* (Fig. 6c,f). The results also showed that (22 ± 3)% and (55 ± 3)% of NECs were associated with *S. flexneri* and the CSF1R or the CSF3R marker, respectively (Fig. 6c,f). These results indicate that the vast majority of infected NECs are neutrophils and macrophages in the lamina propria 24 hpi. It follows that (12 ± 5)% of NECs are associated with *S. flexneri* but do not express CSF1R or CSF3R.

**Fig. 6.**
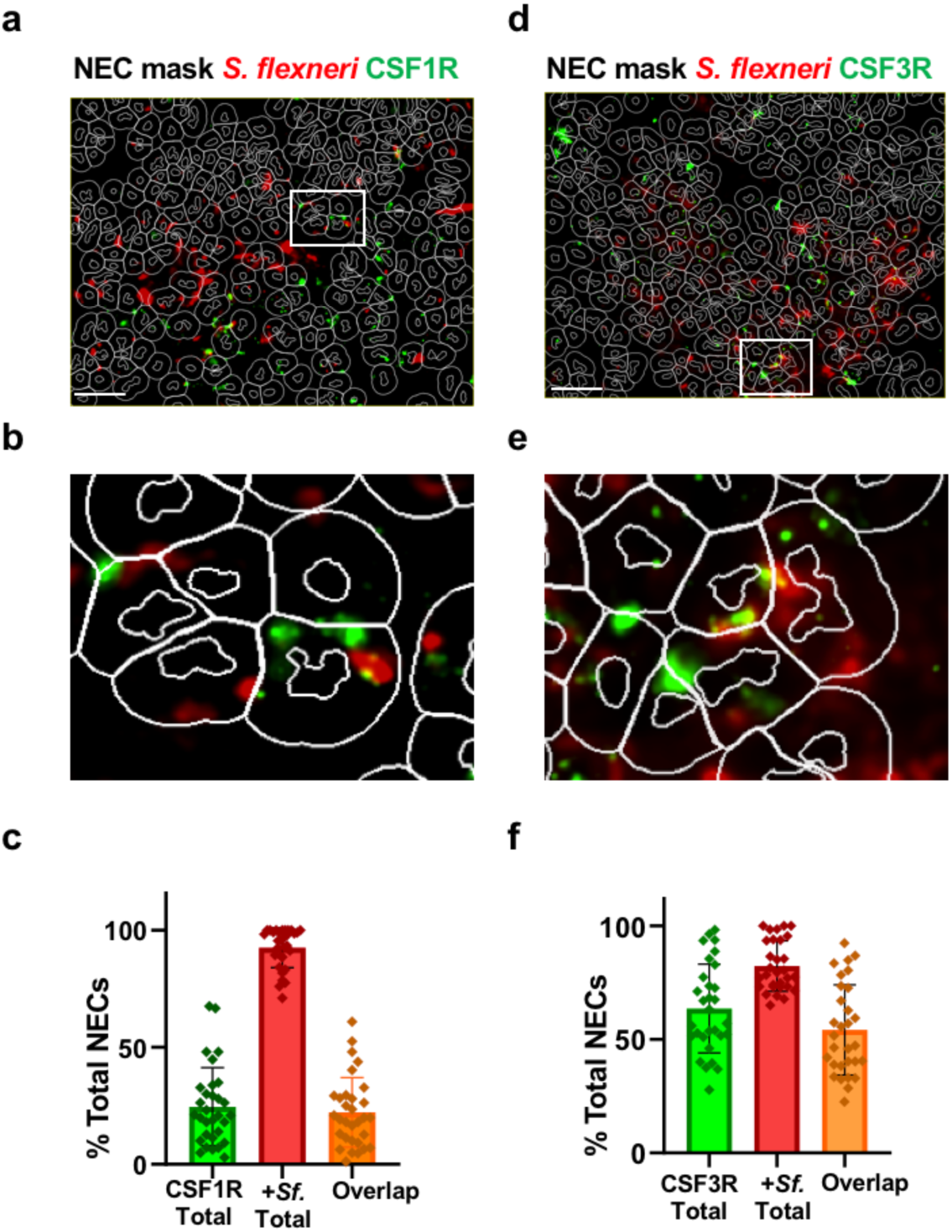
Bacterial association with Neutrophils and Macrophages. (**a-b**) Representative images of colonic mucosa 24 hpi with *S. flexneri* (red) showing NECs (Mask, white) stained for CSF1R (a-b, green) and CSF3R (d-e, green). Scale bar: 100 µm. **(d,f)** Graphs showing the percentage of infected NECs associated with the CSF1R (d) and CSF3R (f) markers as shown in (c,e). Each symbol represents one image; bars show the mean and error bars the SEM (n = 30 images from 2 colons). All measures are expressed as a percentage of DAPI+ nuclei in the same image. CSF1R+ 25 ± 3%; bacteria+ 93 ± 1%; CSF1R+ bacteria+ 22 ± 3%. Cells were segmented on DAPI; CSF1R was detected in the FITC channel and *S. flexneri* by FISH in the TRITC channel, with cells scored positive when subcellular signal was detected within the segmented cell boundary. No statistical comparison is shown between bars, as the co-positive population is a subset of each single-marker population. Observed co-positivity did not differ from that expected under independent association (observed/expected = 0.98). (f) CSF3R and *S. flexneri* association. As above, with CSF3R detected in the FITC channel and *S. flexneri* in the Cy5 channel (n = 30 images from 2 colons). CSF3R+ 65 ± 3%; bacteria+ 86 ± 2%; CSF3R+/ bacteria+ 55 ± 3%. Observed/expected co-positivity = 0.98.

These cells may still belong to the myeloid lineage, such as dendritic cells, or derive from the common lymphoid lineage, such as B and T cells^28^.

## Discussion

In this study, we conducted microscopy analysis of the colonic tissue and invading bacteria in time-resolved infection experiments. In agreement with the notion that shigellosis relies on the ability of the bacteria to invade epithelial cells and spread from cell-to-cell^12^ we observed a temporal correlation between epithelial infection and epithelial fenestration. However, after this early stage of infection, the bacteria transitioned from their primary niche, the epithelial compartment, to their secondary niche, the lamina propria. Transcriptomic analysis led to the identification of various cellular markers, including the macrophage– and neutrophil-specific markers CSF1R and CSF3R, respectively. Importantly, the vast majority of the bacteria were associated with neutrophils or macrophages during the late stage of infection. We discuss the implications of these observations in our understanding of the processes supporting shigellosis.

### Colonic invasion: Direct Invasion of Epithelial Cells

We previously showed that during the early phase of infection in infant rabbits, *S. flexneri* was associated with E-cadherin-positive cells as early as 2 hpi^12^. Here we show that bacteria are uniquely found within the epithelial layer and, importantly, virtually absent from the lamina propria in the distal colon up to 8 hpi. We also detected very few cell death events in the mucosa during this early stage of infection. The current model of shigellosis posits that the primary niche of infection are macrophages in the lamina propria. This model is based on infection experiments conducted in the small intestine of adult rabbits^14^, in which *S. flexneri* transcytoses via M cells from the lumen to the lamina propria, where it encounters, infects and kills macrophages in less than 4 hours^16^. However, the primary site of symptomatic infection is the colon in humans^29,30^, and M cells are very sparse in the colonic mucosa^31^. Therefore, the infant rabbit model of shigellosis may offer a novel appreciation of the route of invasion in the colonic environment. Given that *S. flexneri* infects polarized cells via the basal-lateral pole of epithelial cells in tissue culture systems^13,32^, how bacteria circumvent the physical barrier represented by tight junctions for efficient infection *in vivo* and whether immune cell infiltration may assist in the process remains to be determined.

### Decrease in marker representation: Epithelium Collapse

*S. flexneri* infection in humans, non-human primates, guinea pigs, and rabbits leads to severe ulceration of the colonic mucosa^4,5,33–35^. Accordingly, the time course analysis conducted here in infant rabbits revealed an increasing loss of E-cadherin-positive cells in infected samples, showing a continuous loss of epithelial integrity as infection progresses. In support of this notion, global gene expression profiles of the corresponding samples showed a decrease in representation of genes associated with various epithelial cell markers (Supplementary Table 1). These genes include LGR5, a stem cell marker^36^, CLCA1, a goblet cell marker^37^, LYZ, a Paneth cell marker^38^, and SLC26A3, a differentiated colonocyte marker^39^. Depletion of a stem cell marker may lead to defects in the regeneration program^40^. Depletion of goblet cells may have an impact on mucus-supported barrier functions^41^. Depletion of differentiated enterocytes may impact the nutrient acquisition and barrier functions of the epithelium, as reflected by animal weight loss^12^. It is common knowledge that shigellosis is an inflammatory disease in which neutrophil infiltration is chiefly responsible for the tissue damage inflicted to the epithelial tissue upon *S. flexneri* infection^19,21^. However, we have previously demonstrated the critical role of bacterial cell-to-cell spread in challenging the epithelial integrity in infant rabbits^12^. How bacterial dissemination in the colonic tissue affects epithelial integrity and what exact role neutrophils play in that process remains to be determined.

### Increase in marker representation: Cytokine/Chemokine signaling and Immune Cell Recruitment

The early immune response to Shigellosis has been characterized in human colonic samples and in adult rabbit intestine, showing massive infiltration of myeloid cells^19,42,43^. *S. flexneri* infection also led to infiltration of monocytes and neutrophils in infant rabbits^12^. In agreement with these results, the transcriptomics analysis revealed a dominant innate myeloid inflammatory signature with strong evidence of neutrophil and monocyte/macrophage infiltration (Supplementary Table 1). This is reflected through deconvolution of the RNA-Seq results leading to the identification of cellular sub-types and modules (Fig. 4). The approach revealed specific markers for neutrophils (CSF3R, CXCR1/2, CEBPE, LCN2, S100A8/9, MS4A3, TCN1, CAMP, LTF, CRISP3, PGLYRP1)^44^, monocytes (CD14, ITGAM, TYROBP, FCER1G) and macrophages (CSF1R, CD163, MRC1, MARCO, MSR1, APOC1, VSIG4, C1R, C1S)^45^. The infiltration of these immune cells is potentially supported by adhesion factors, such as ICAM1 and SELP on activated endothelial cells and SELL and ITGB2 on immune cells^46^. Circulating immune cells may respond to chemotactic signaling originally emanating from infected epithelial cells in the form of chemokines such as CXCL8 and CCL2, whose production has been demonstrated in human biopsies in response to shigellosis^47^. CXCL8 and CCL2 signal through their cognate receptors, CXCL1/2 and CCR2 on neutrophils and macrophages, respectively. Infiltrating cells are exposed to various inflammatory cytokines shaping their transcriptional profiles (TNF, IL1B, IL6,etc…), whose expression has also been demonstrated in shigellosis^47^. In addition to cell types and cognate signaling, cellular deconvolution allowed speculation on cellular functions. Numerous identified markers qualify as neutrophil-specific effectors supporting antimicrobial activity (LCN2, LTF, CAMP, PGLYRP1), tissue remodeling (MMP8, MMP9), ROS production (NCF2, CYBB/NOX2), phagocytosis (ITGAM, FCER1G, C5AR1)^48^. Macrophage-specific markers support phagocytosis of opsonized bacteria (FCGR1A / FCGR3A, ITGAM) and bacterial particles (MARCO, MRC1, CD36), and phagosome maturation (CTSs, LAMPs, ATP6V1/ATP6V0)^49^. In summary, gene expression profiles support a massive recruitment of immune cells in the form of neutrophils and macrophages, with potential functions in bacterial clearance.

### Colonic invasion: Late Invasion of Immune Cells

A critical development of this study is the demonstration that *S. flexneri* first invades and spreads in epithelial cells, and then disseminates to the lamina propria where it interacts with immune cells. This is in contrast with the current model of shigellosis in which the first niche of infection is the macrophages (Fig. 7). Thus, in the small intestine of adult rabbits, the first events of inflammation essentially occur in immune cells in the lamina propria, whereas in the colon of infant rabbits, epithelial cells are likely to be the cellular sub-type initiating inflammation. Future experiments will be required to determine the exact contribution of primary inflammatory events occurring in epithelial cells and secondary downstream events leading to (i) activation of resident cells in the lamina propria and (ii) the recruitment of circulating immune cells (Fig. 7).

The use of cell type-specific markers demonstrated that, during the late phase of infection, bacteria exist in association with neutrophils and macrophages. The association with neutrophils is not surprising as these phagocytes have been proposed to be the main immune cells supporting bacterial clearance, albeit at the cost of host tissue damage^19^. However, we showed that bacterial burden dramatically increased during the late phase of infection, when virtually all neutrophils were associated with bacteria. It is thus unclear where and how the bacterial population expanded in the tissue and the exact role played by neutrophils in bacterial clearance remains to be clarified.

Recently, a role for macrophages has been suggested in controlling infection in an immunocompromised mouse model of shigellosis^50^. The proposed mechanisms involved a regulatory loop supported by macrophage production of IL12 driving IFN-γ-mediated restriction of bacterial growth in epithelial cells. Whether bacteria associated with macrophages or neutrophils was not investigated in this study. Given that, similar to the situation observed with neutrophils, the late phase of infection in the rabbit model corresponds to very high bacterial burden in macrophages, whether the observation of *S. flexneri* presence within immune cells corresponds to bacterial clearance, or may represent a novel strategy of colonization remains to be investigated.

**Fig. 6.**
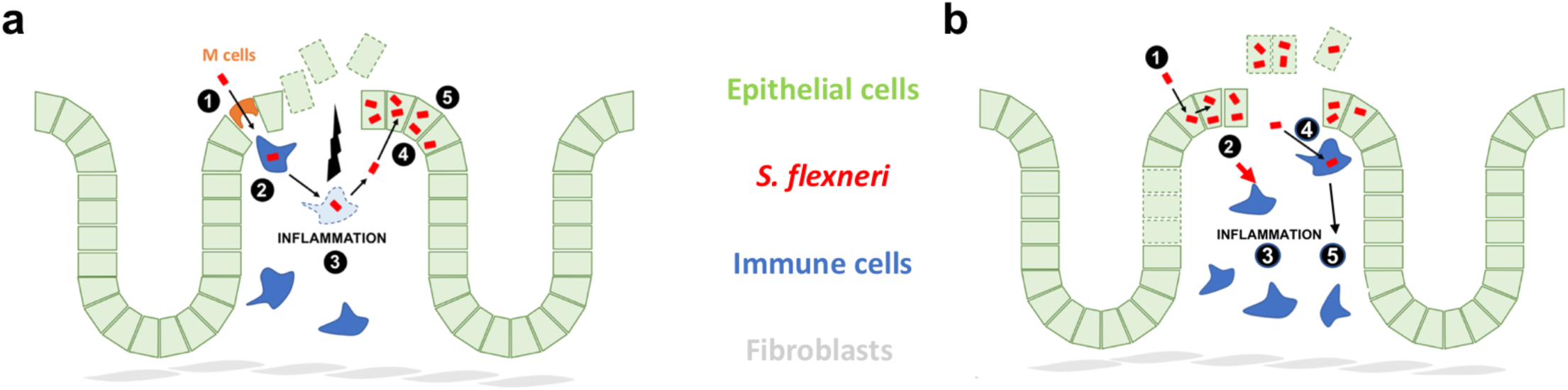
Current and novel models of shigellosis. **(a) Current model of shigellosis.** The model is derived from studies conducted in the adult rabbit small intestine. (**1**) *S. flexneri* transcytoses through M cells. (**2**) *S. flexneri* encounters and invades macrophages. (**3**) Infected macrophages are activated and killed, which triggers inflammation and tissue destruction. (**4**) Bacteria released from dead macrophages invade epithelial cells through their basal side. (**5**) *S. flexneri* establishes its major niche in epithelial cells through cell-to-cell spread. **(b) Novel model of shigellosis.** The model is derived from studies conducted in the infant rabbit colon. (**1**) *S. flexneri* invades epithelial cells through their apical-lateral side, grows, and spreads from cell-to-cell. (**2**) Infected epithelial cells produce inflammatory cytokines/chemokines, and bacterial spread from cell to cell leads to epithelial cell fenestration. (**3**) Immune signaling and tissue damage lead to immune cell recruitment and severe inflammation. (**4**) Bacteria gain access to the lamina propria and associate with immune cells, which may lead to increased inflammation and cell death, but also transient colonization. (**5**) Ultimately, immune cells resolve the infection process through signaling coordination, bacterial killing, tissue remodeling, and debris removal.

## Material and Methods

### Bacterial Strains

The wild-type *S. flexneri* strain used in this study was serotype 2a 2457T and was cultured in tryptic soy broth (TSB) (Sigma-Aldrich, Cat. No. 22902) medium at 37 °C.

### Animal Procedures

Pregnant female New Zealand White rabbits were obtained from a commercial breeding company (Charles River). Newborns were isolated after birth and kept in a 30°C incubator for the duration of the experiment. At the time of feeding, does were dosed with 0.3ml oxytocin (Bimeda, Cat. No. 069241) and 0.5ml Acepromazine injectable solution (Covetrus, Cat. No. 003845), infant rabbits were fed on the tranquilized does and put back into the incubator.

For infection of 10-15-day old kits, *S. flexneri* was grown for 12.5 hours at 37°C in 5mL TSB per animal on a rotating wheel. The cultures were pelleted and resuspended in 200 µL PBS (Gibco Cat. No. 14190144) per animal prior to inoculation. Infant rabbits were anesthetized in a chamber with 5% isoflurane in 5 L/min oxygen before rectal inoculation with 200 ul of bacterial resuspension using feeding tubes. Post-inoculation, the infant rabbits were weighed daily pre– and post-feeding and observed for clinical signs of illness.

All in vivo experiments described here were reviewed and approved by the University of Virginia Institutional Biosafety Committee and the Institutional Animal and Care and Use Committee. Care of the does and infant rabbits adhered to standard operating procedures developed in coordination with the veterinary and animal care staff of the Center for Comparative Medicine at the University of Virginia.

### Histology

At the appropriate time points, infant rabbits were euthanized by CO2 (Praxair, Cat. No. CD50) inhalation and 0.1mL of euthasol (Virbac, Cat. No. NDC 51311-050-01) and the colon was harvested. The colons were rinsed and flushed with modified bouin’s fixative solution using equal parts of ethanol (Fisher Cat. No. 04-355-223) and water with 5% glacial acetic acid (Fisher, Cat. No. A38-500), cut longitudinally, arranged in cassettes as swiss rolls, and placed into neutral-buffered 10% formalin (Fisher, Cat. No. SF100-4) for paraffin sections. After 48 h, samples were moved to 70% EtOH for storage before loading onto a tissue processor for dehydration and paraffin infiltration. After manual embedding into a paraffin block, paraffin sections were cut at 5 µm on UVA Histology Core.

### Histopathology Analysis

Paraffin sections of colonic samples were stained with hematoxylin and eosin at UVA Histology Core. Slides were scanned using an Aperio Scanscope CS5 scanner, and snapshots at 40x were taken of the entire colonic tissue. Using the lasso tool in Aperio ImageScope 12.3.3, red blood cells seen in the mucosa and sub-mucosa were outlined (Fig. 2), and the blood area was measured.

### Immunofluorescence Microscopy

Paraffin sections were rehydrated in a graded series of xylene (Fisher, Cat. No. X3F), 100% ethanol, 95% ethanol, 70% ethanol, and water. Antigen retrieval was performed in an Instant Pot using pre-heated citric acid-based buffer (Vector Laboratories, Cat. No. H-3300): buffer was pre-heated on the High Sauté setting for 10 min, slides were loaded onto a slide holder and submerged, and the Instant Pot was run on High-Pressure Cooker mode for 1 min. Pressure was released using the quick-release method, and slides were transferred to water for 15 min to cool, followed by three 3-min washes in PBS. Slides were permeabilized in 1% Triton X-100 for 10 min at room temperature and washed three times in PBS (3 min each). Slides were blocked for 1 h at room temperature in 2% normal goat serum (Vector Labs, Cat. No. S-1000-20) and 5% bovine serum albumin (AmericanBio, Cat. No. AB01088-00100) in PBS, followed by a single 3-min PBS wash.

Primary antibodies were diluted 1:100 in blocking buffer and incubated overnight at 4°C. For fenestration and bacterial burden analyses, sections were stained with anti-E-cadherin (BD Biosciences, Cat. No. 610181) and anti-*Shigella* sp. (ViroStat, Cat. No. 0901). For cell-type infection analyses, sections were stained with anti-E-cadherin and anti-*Shigella* sp. (ViroStat, Cat. No. 0901). Slides were washed three times in PBS (3 min each), then incubated with appropriate secondary antibodies, including Goat anti-Mouse Alexa Fluor™ 568 (ThermoFisher, Cat No. A-11004) and Goat anti-Rabbit IgG (H+L) Highly Cross-Adsorbed Secondary Antibody after diluting it at a ratio of 1:500 in blocking buffer for 2 h at room temperature. Coverslips were mounted with ProLong Gold Antifade Mountant (Thermo Fisher Scientific, Cat. No. P36930).

### Dual RNAscope Immunofluorescence

Paraffin sections were deparaffinized and dehydrated in the following sequence: xylene, 100% ethanol, and hydrogen peroxide. Antigen retrieval was performed at 95-100 °C for 15 min using boiling 1x RNAscope Target Retrieval Reagent (Advanced Cell Diagnostics, Cat. No. 322000). Slides were rinsed in deionized water for 15 seconds and washed in 100% ethanol for 3 min.

Slides were permeabilized with RNAscope Protease Plus for 30 min at 40°C within the RNAscope HybEZ II Hybridization System. Slides were washed in RNAscope Wash Buffer (Advanced Cell Diagnostics, Cat. No. 310091). Probes were hybridized on the slides for 2 hours at 40°C within the RNAscope HybEZ II System. Slides were washed in RNAscope Wash Buffer. Slides were incubated within the RNAscope HybEZ II System using the RNAscope Multiplex Fluorescent Detection Reagents v2 kit (Advanced Cell Diagnostics, Cat. No. 323110) in order at 40°C: AMP 1 (Advanced Cell Diagnostics, Cat. No. 323101) for 30 minutes, AMP 2 (Advanced Cell Diagnostics, Cat. No. 323102) for 30 minutes, and AMP 3 (Advanced Cell Diagnostics, Cat. No. 323103) for 15 minutes.

HRP-C1 (Advanced Cell Diagnostics, Cat. No. 323104) for 15 minutes, Opal 690 Reagent (Akoya Biosciences, Cat. No. OP-001006) at 1:1000 within RNAscope Multiplex TSA Buffer (Advanced Cell Diagnostics, 322809) for 30 minutes, and HRP blocker (Advanced Cell Diagnostics, Cat. No. 323107) for 15 minutes, HRP-C2 (Advanced Cell Diagnostics, Cat. No. 323104) for 15 minutes, Opal 520 Reagent (Akoya Biosciences, Cat. No. OP-001006) at 1:1000 within RNAscope Multiplex TSA Buffer (Advanced Cell Diagnostics, Cat. No. 322809) for 30 minutes, and HRP blocker (Advanced Cell Diagnostics, Cat. No. 323107) for 15 minutes, HRP-C3 (Advanced Cell Diagnostics, Cat. No. 323104) for 15 minutes, Opal 590 Reagent (Akoya Biosciences, Cat. No. OP-001006) at 1:1000 within RNAscope Multiplex TSA Buffer (Advanced Cell Diagnostics, Cat. No. 322809) for 30 minutes, and HRP blocker (Advanced Cell Diagnostics, Cat. No. 323107) for 15 minutes. After each incubation step, slides were washed in RNAscope Wash Buffer for 5 minutes with gentle agitation.

Immediately following the RNAscope procedure, slides were blocked with 2% normal goat serum in 5% bovine serum albumin (AmericanBio, Cat. No. AB01088-00100) PBS for 1 h at room temperature. Primary antibodies were diluted 1:100 in blocking buffer (E-cadherin, BD Biosciences Cat. No. 610181) and incubated overnight at RT. Secondary antibodies (goat anti-mouse Alexa Fluor Pacific Blue) were diluted 1:500 in blocking buffer and incubated for 2 h at room temperature. Slides were counterstained using RNAscope Multiplex FL v2 DAPI (Advanced Cell Diagnostics, Cat. No. 323108) for 1 minute. Coverslips were mounted using ProLong Gold Antifade Mountant. Slides were imaged using a Nikon TE2000 microscope equipped for automated multi-color imaging, including a motorized stage and filter wheels, a Hamamatsu Orca ER Digital CCD Camera and piezo-driven 10X and 60X objectives. All Images were processed with the Qupath image analysis software V7^24^, and the counts for the RNA probes used were performed using the reported protocols^51^.

### Transcriptomic Analysis

#### (i) RNA isolation

Post-RNase inactivation, the extracted colon was subjected to RNA isolation using TRIzol (Life Tech, Cat. No. 15596018). Tissues were homogenized, followed by precipitation using isopropanol and ethanol (Yum et al., 2019). The isolated RNA was resuspended in RNase-free water and used for transcriptomic analysis.

#### (ii) Illumina library construction

Steps involving library preparation and RNA sequencing were outsourced to Novogene (CA, USA). Concisely, the library construction involved sequential steps of RNA quantification and qualification, mRNA enrichment using Ribo-Zero rRNA removal kit specific for bacterial rRNA (Illumina, MRZMB126), cDNA synthesis, end repair and adaptor ligation, size selection of fragments, and PCR followed by quality check. The nucleic acid quality check steps included NanoDrop and agarose gel electrophoresis; the extracted RNA was assessed using a 2100 Bioanalyzer (Agilent Technologies) before sequencing.

#### (iii) RNA sequencing

NEB Next Ultra II RNA Library Prep kit was used by following the manufacturer’s instructions for Illumina (BioLabs, New England, MA, USA). The first step comprises strand-specific library synthesis to replace the dTTPs with dUTPs while synthesizing the second-strand cDNA. Resulting overhangs were converted to blunt ends, after which adenylation of 3′ ends was performed, followed by adapter ligation using RNA 5′ and RNA 3′ adapters provided in the kit. An insert size check was performed during quality check steps, and a library concentration of 1.5 ng/μL was used for sequencing. Paired-end sequencing was performed using libraries thus generated on the Illumina platform, and the next set of steps comprised cluster growth and sequencing, image acquisition, and base-calling.

#### (iv) Differential Gene Expression Analysis

The raw reads were subjected to a typical RNASeq analysis as described earlier^52^. The rabbit genome sequences and associated transcriptomic files available in NCBI (Assembly Name: OryCun2.0 INSDC / GenBank Assembly ID: GCA_000003625.1) and the Ensembl database (https://useast.ensembl.org/index.html accessed in September 2022) were used to count gene hits^53^. An annotation file was used to generate fold-change calculations and transcripts per million (TPM) values using Kallisto. Normalized counts were retrieved after addressing variation in the size of data sets and varying gene lengths. Gene assignments for estimated counts less than 100 in either one of the samples being compared were removed. The resulting files were used to calculate fold change and log2 change values using DESeq2^54^. The differentially expressed genes (DEGs) were tabulated based on these calculations.

#### (v) C8 analysis

Gene expression was quantified from RNA-sequencing reads and normalized using the regularized-logarithm (rlog) transformation in DESeq2. For visualization of individual cell types C8 database was used with recommended run parameters^55^.

### Immunofluorescence Image Acquisition and Quantification

Slides were imaged using NIS-Elements software (version AR6.02.03; Nikon Instruments Inc., Avon, Massachusetts USA) controlling a Nikon TE2000 microscope equipped with a Hamamatsu Orca-ER camera (Hamamatsu Photonics, Hamamatsu City, Japan). For fenestration and bacterial burden analyses, whole-colon imaging was performed (Supplementary Fig. 1 and Fig. 2). Images were processed in Leica ImageScope and QuPath for quantification of epithelial fenestration (Fig. 2), classification of cell types (ECs vs. NECs), nuclear morphology sorting, and enumeration of marker-positive cells and co-localization of signals.

Colonic tissue sections from PBS and *S. flexneri*-infected infant rabbits were stained for E-cadherin, *S. flexneri*, CSF1R, CSF3R, LCN2 and MARCO (Advanced Cell Diagnostics, Oc-CSF1R-C2: Cat. No. 857991; Oc-CSF3R-C1: Cat. No. 1081031-C1; Oc-LCN2-C2: Cat. No. 857241; Oc-MARCO-C1: Cat. No. 847598) as indicated in the corresponding figures. Stained sections were imaged, and cells were computationally segmented and classified based on fluorescence channel positivity into single-positive, double-positive, and double-negative populations for each marker combination assessed.

#### (i) Bacteria and Marker Co-localization Analysis

To assess co-localization of luminal bacteria with CSF1R or CSF3R cells, segmented cell counts for each field of view were grouped into three mutually exclusive categories: marker-positive only, bacteria-positive only, and double-positive: marker-positive & bacteria-positive (co-localized/overlap). Cells positive for bacteria or marker were assigned to the corresponding bacteria-only, marker-only, or overlap category, and values were expressed as raw cell counts and as a percentage of the sum of the three categories per colon. Data were derived from 2 biological replicates (animals), each with 2 technical replicates (colon sections/images), for a total of 4 measurements per marker.

#### (ii) Epithelial Fenestration and Blood Area Quantification

Hematoxylin and eosin-stained colonic sections from mock (PBS) and S. flexneri-infected animals were assessed for epithelial fenestration and luminal/mucosal blood area. Percent fenestration was calculated as the fenestrated length relative to the total measured colon length per sample. Blood area was measured across the total length of infected colon tissue at each time point (4, 8, and 24 h post-infection).

#### (iii) Bacterial Burden Quantification

Bacterial burden was quantified per tissue sample across mock and infected conditions and time points (4, 8, and 24 h post-infection) and separately compared between E-cadherin-positive epithelial cells (ECs) and non-E-cadherin (NEC) cell populations at each time point. Objects equal to the number of bacteria in images acquired at 10X resolution from colon were counted using thresholding and subcellular spot detection in QuPath. Densely packed bacteria that could not be resolved as individual objects were assigned an estimated spot count by dividing the detected region area by the expected single-bacterium area (1 µm², minimum 0.5, maximum 2). Values therefore represent estimated bacterial spot counts summed across constant images per colon, rather than absolute bacterial numbers.

#### (iv) Cell Death Assay

Apoptotic cells were identified by TUNEL staining and quantified in QuPath using the same segmentation approach. TUNEL-positive cells are reported as a percentage of (total nuclei per field/total cells in the compartment), scored separately in EC and NEC compartments within the same fields.

#### (v) Statistical Analysis

All statistical analyses were performed in GraphPad Prism V10 (for Windows, Boston, Massachusetts, USA). Tests were selected on the basis of the distribution of each dataset, and all p-values are two-tailed and multiplicity-adjusted within the family stated. Comparisons across two independent factors (e.g., condition × time, or cell type × time) were analyzed by two-way ANOVA, Tukey’s multiple comparisons test applied post hoc or t-test with Holm’s correction, as indicated in individual figure legends. Comparisons across three or more groups within a single factor were analyzed by ordinary one-way ANOVA with Tukey’s or Šídák’s multiple comparisons test; where variance was non-normally distributed, data were log-transformed prior to analysis.

Homogeneity of variance was assessed using Brown-Forsythe and Bartlett’s tests. For bacterial burden, within-group variance scaled with the mean, groups were compared by Brown-Forsythe and Welch ANOVA with the Games-Howell post-test. Data are presented as mean ± SEM or mean ± SD as indicated in each figure legend, with individual biological replicates shown as single data points. A P value < 0.05 was considered statistically significant (ns, not significant; \**p*< 0.05; \*\**p*< 0.01; \*\*\**p*< 0.001; \*\*\*\**p*< 0.0001).

## Data Availability

The RNASeq data generated from infected and control samples used in this study are available upon request.

## Acknowledgements

We thank all the members of the Agaisse laboratory for discussions on the project. We thank Mahua Mandal for coordinating the laboratory rabbit colony. We thank Alice Kweon and Zackary Lifschin for nursing infant rabbits. We thank the Staff of the LiSA Animal Facility for veterinary assistance. We thank Sheri Vanhoose and the UVA Research Histology Core for preparing paraffin sections.

## Author contributions

NBJ: investigation, methodology, formal analysis, visualization, writing, bioinformatics and image analysis.

BAD: investigation, methodology, formal analysis, visualization, writing, image analysis.

LKY: investigation, methodology.

HFA: conceptualization, supervision, funding acquisition, project administration, writing: review & editing. All authors read and approved the final manuscript.

## Competing interest

The authors declare that no competing interests exist.

## Funding

This work was supported by the National Institutes of Health grant R01AI073904, R01AI179778, and R21AI149384 (H.A.).

## Supplementary Files

### Supplementary Figures

**Supplementary Fig 1. Image analysis of the infected colonic mucosa**.

**Supplementary Fig. 2. Cellular infection in Lamina Propria**.

### Supplementary Tables

**Supplementary Table 1. Annotation of DeSeq2 analysis using the C8 data base**.

**Supplementary Fig. 1.**
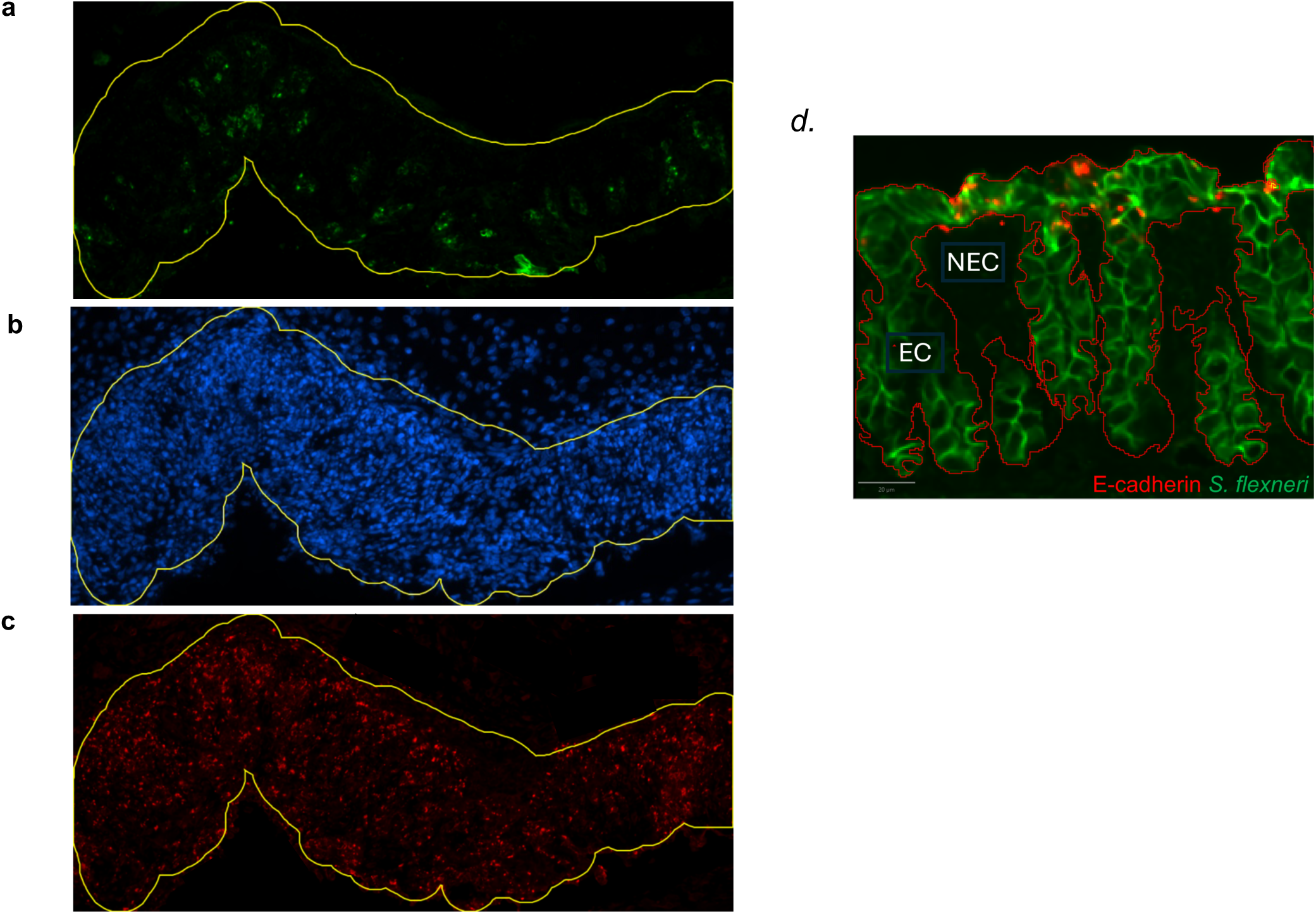
Image analysis methods in the infant rabbit model of bacillary dysentery. Representative fluorescence images illustrating the bacterial quantification pipeline applied to infected colon tissue collected at 24 hpi. **(a)** DAPI (blue) nuclear staining, which is combined with **(b)** E-cadherin (ECAD; green) staining of the intestinal epithelium, which is largely degraded by 24 hpi residual ECAD signal to delineate and create a mask of the mucosal region and measure its corresponding area. **(c)** Bacterial signal (red) in a separate channel. **(d)** Compartment based estimation of bacterial counts within EC and NEC region of Mucosa in infections across time.

**Supplementary Fig. 2.**
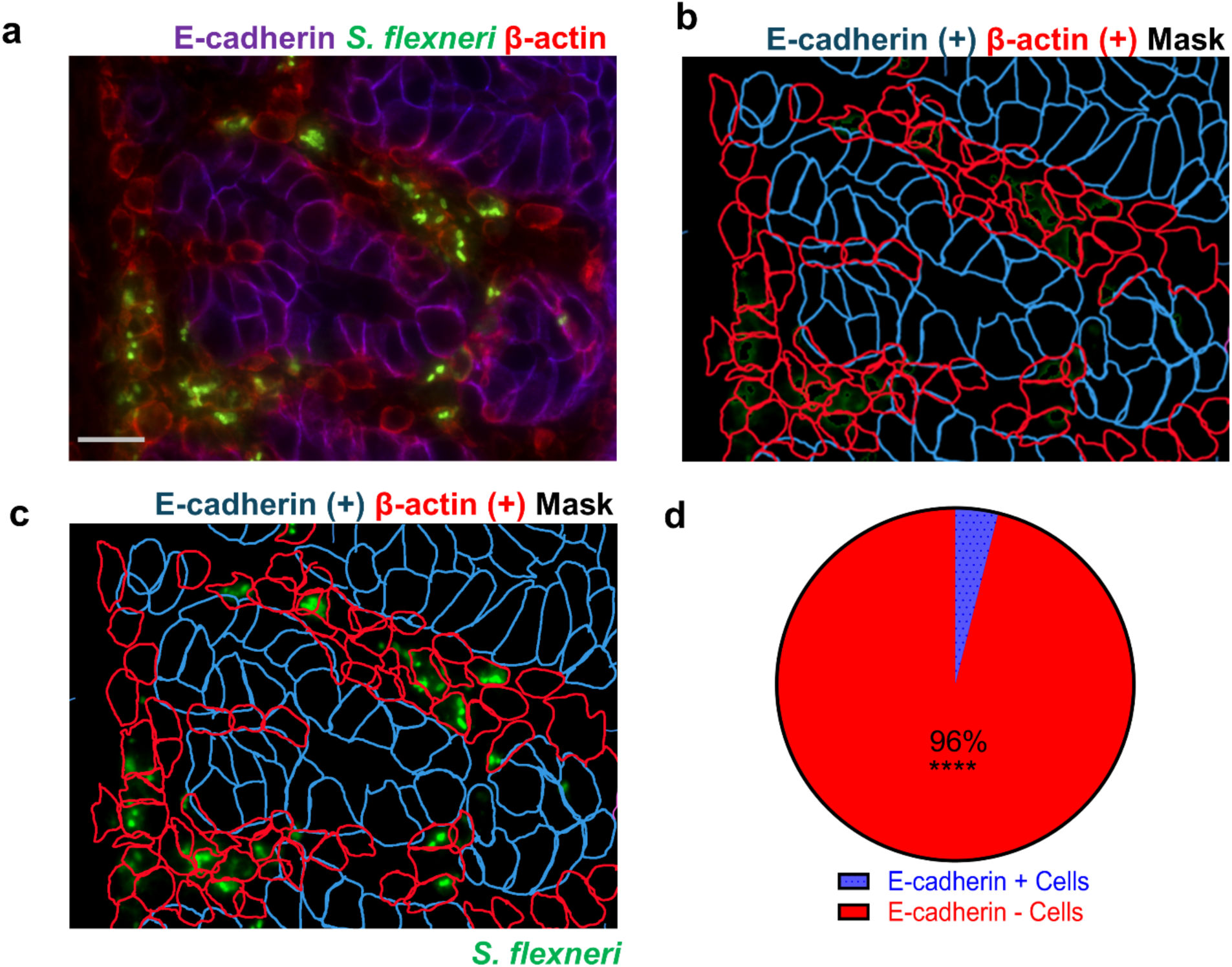
Cellular infection in the lamina propria. **(a)** Representative image of a colonic section at 24 hpi, immuno-stained for E-cadherin (purple), *S. flexneri* (green) and β-actin (red). Scale bar, 50 µm. **(b)** Segmentation mask distinguishing E-cadherin-positive cells (blue outlines) from E-cadherin-negative, β-actin-positive cells (red outlines). **(c)** Overlay of the mask in (b) with the *S. flexneri* channel (green). **(d)** Percentage of infected cells that were E-cadherin-positive (epithelial, blue) or E-cadherin-negative (lamina propria, red), quantified from (c) Percentage of infected cells that were E-cadherin-positive (epithelial) or E-cadherin-negative (lamina propria), quantified from (**c**). n=20 imaging fields; individual fields were treated as replicates. Statistics: Wilcoxon matched-pairs signed-rank test, median difference 47.0%, ****P < 0.0001.

**Supplementary Table 1.** Annotation of rlog normalized reads analysed using the C8 data base. Conditions: mock-treated samples (PBS, n=5) and, infected samples (WT n=5). Values are on the rlog scale.

| Name | 24PBS1 | 8PBS2 | 24PBS2 | 8PBS3 | 24PBS3 | 24WT1 | 24WT2 | 24WT4 | 24WT5 | 24WT3 |
| --- | --- | --- | --- | --- | --- | --- | --- | --- | --- | --- |
| LGR5 | 10.4808 | 10.4398 | 10.5307 | 11.0022 | 10.6141 | 9.9503 | 9.3003 | 9.4542 | 9.6573 | 9.6451 |
| AXIN2 | 10.0094 | 10.0258 | 10.0071 | 9.6753 | 10.0236 | 9.5749 | 9.7211 | 9.519 | 9.3698 | 9.6868 |
| PLAGL2 | 9.7223 | 9.9553 | 9.832 | 9.9407 | 9.9892 | 9.1239 | 9.3287 | 9.2564 | 9.4012 | 9.2411 |
| SOX9 | 9.6354 | 9.6661 | 9.5036 | 9.8094 | 9.4281 | 9.3159 | 8.8733 | 9.3796 | 9.2537 | 9.0337 |
| LYZ | 19.0603 | 18.3632 | 19.1155 | 18.5928 | 18.9268 | 17.6449 | 16.9699 | 17.3936 | 17.9874 | 17.6669 |
| WNT3 | 1.3204 | 1.4754 | 1.4038 | 1.5367 | 1.7135 | 1.3274 | 1.3986 | 1.527 | 1.3208 | 1.3601 |
| ATOH1 | 10.1693 | 10.0738 | 10.0098 | 10.2349 | 9.8475 | 9.3139 | 8.3507 | 9.1681 | 9.2782 | 9.2041 |
| B3GNT6 | 9.7459 | 10.1132 | 9.6737 | 10.1756 | 9.8122 | 9.6049 | 8.9396 | 9.5142 | 9.8305 | 9.2127 |
| CLCA1 | 12.9814 | 13.3616 | 13.1456 | 13.6118 | 13.0618 | 12.0627 | 11.3999 | 11.9722 | 12.5569 | 11.8426 |
| SLC37A1 | 10.6453 | 10.7258 | 10.5639 | 10.658 | 10.5388 | 10.3669 | 10.0968 | 10.3736 | 10.1697 | 10.1882 |
| CLDN8 | 11.515 | 11.4798 | 11.1852 | 11.6913 | 11.617 | 10.213 | 9.3919 | 10.356 | 10.5537 | 10.3019 |
| AADAC | 9.6469 | 9.5418 | 9.6138 | 9.5266 | 9.5305 | 8.2829 | 7.9037 | 8.3994 | 9.0206 | 8.3522 |
| SLC26A3 | 10.6964 | 11.1913 | 11.681 | 11.178 | 11.5254 | 10.1882 | 8.3306 | 8.8989 | 9.7715 | 9.4657 |
| RNF186 | 8.9916 | 8.464 | 8.7559 | 8.4351 | 8.9163 | 6.8108 | 6.1156 | 6.9427 | 7.1473 | 7.2974 |
| CSF3R | 6.4618 | 6.2879 | 6.0108 | 6.5777 | 6.1086 | 8.8447 | 10.2148 | 10.3696 | 10.0846 | 9.1945 |
| LCN2 | 7.774 | 7.792 | 7.7663 | 8.1684 | 7.7528 | 11.8265 | 13.0889 | 13.3101 | 12.672 | 12.0071 |
| LTF | 6.7501 | 6.7187 | 6.9455 | 6.8608 | 6.8812 | 9.5811 | 10.3388 | 9.9578 | 10.2968 | 10.3442 |
| CAMP | 6.4612 | 6.5325 | 6.41 | 6.4658 | 6.4045 | 9.9865 | 11.4167 | 11.4401 | 11.19 | 10.5415 |
| MMP9 | 5.8787 | 6.6783 | 5.9773 | 6.2661 | 5.654 | 8.9075 | 9.7074 | 10.1244 | 9.8179 | 8.4893 |
| CD14 | 7.6347 | 7.532 | 7.2952 | 7.5087 | 7.2253 | 9.6041 | 10.3711 | 10.2643 | 10.1178 | 9.6819 |
| OLR1 | 5.6459 | 5.5868 | 5.8679 | 5.9746 | 5.6632 | 8.1038 | 8.8778 | 9.2557 | 8.6933 | 8.4287 |
| CD68 | 9.8989 | 10.1939 | 9.6091 | 9.7433 | 9.8925 | 10.1474 | 10.2514 | 10.8252 | 10.3318 | 10.158 |
| CCL2 | 6.4282 | 6.4269 | 6.6739 | 7.2035 | 6.472 | 10.8677 | 11.1709 | 10.644 | 8.8962 | 10.0971 |
| CSF1R | 9.1063 | 9.0029 | 8.7587 | 9.0239 | 8.7481 | 9.2358 | 9.8502 | 10.1657 | 10.3145 | 9.2795 |
| TNF | 5.0378 | 5.0912 | 5.0742 | 5.1913 | 4.8108 | 7.7998 | 8.0709 | 7.9443 | 7.0864 | 7.2698 |
| IL1B | 6.9959 | 6.8756 | 6.9364 | 7.1733 | 6.8522 | 10.9771 | 11.8773 | 11.5888 | 10.4826 | 11.0902 |
| CXCL10 | 7.6931 | 7.3148 | 7.1403 | 7.8997 | 6.645 | 9.9543 | 10.3107 | 10.2425 | 9.272 | 9.253 |
| MRC1 | 9.8329 | 9.5812 | 9.7616 | 9.7572 | 9.6296 | 11.0126 | 11.8127 | 11.3195 | 12.3747 | 11.3546 |
| MARCO | 4.945 | 4.9319 | 4.8673 | 5.0276 | 4.7157 | 6.3611 | 7.6036 | 7.3935 | 7.3041 | 5.7607 |
| LYVE1 | 7.8789 | 7.7222 | 7.8551 | 7.8972 | 8.076 | 8.3847 | 8.6423 | 8.4843 | 8.5811 | 8.481 |
| CD86 | 5.8482 | 5.5461 | 5.435 | 5.3701 | 5.0738 | 5.5529 | 5.7217 | 5.6011 | 5.4564 | 5.3651 |
| IL6 | 6.2741 | 6.1931 | 6.1349 | 6.2464 | 6.1253 | 11.3665 | 11.4421 | 10.7825 | 8.7439 | 10.7129 |
| CD163 | 8.1138 | 8.0266 | 7.9764 | 8.2299 | 8.0475 | 9.4916 | 9.8978 | 9.9918 | 10.0415 | 9.4906 |
| ARG1 | 5.4575 | 5.1812 | 5.4172 | 5.7678 | 5.2045 | 7.5947 | 8.6825 | 8.6452 | 8.6212 | 7.7368 |
| CD31 | 10.1979 | 10.3201 | 10.1531 | 10.3668 | 10.1745 | 11.1781 | 11.6335 | 11.6653 | 11.6027 | 11.0925 |
| F3 | 8.7844 | 8.4195 | 8.4348 | 8.5193 | 8.5817 | 11.2963 | 11.2388 | 11.9931 | 10.6581 | 11.2161 |

## References

1. Centers for Disease Control and Prevention. https://www.cdc.gov/shigella/about/index.html (2024).

2 Khalil, I. A. et al. Morbidity and mortality due to shigella and enterotoxigenic Escherichia coli diarrhoea: the Global Burden of Disease Study 1990-2016. Lancet Infect Dis, doi:10.1016/S1473-3099(18)30475-4 (2018).

3 Baker, S. & Scott, T. A. Antimicrobial-resistant Shigella: where do we go next? Nat Rev Microbiol 21, 409–410, doi:10.1038/s41579-023-00906-1 (2023).

4 Takeuchi, A., Formal, S. B. & Sprinz, H. Experimental acute colitis in the Rhesus monkey following peroral infection with Shigella flexneri. An electron microscope study. Am J Pathol 52, 503–529 (1968).

5 Labrec, E. H., Schneider, H., Magnani, T. J. & Formal, S. B. Epithelial Cell Penetration as an Essential Step in the Pathogenesis of Bacillary Dysentery. J Bacteriol 88, 1503–1518 (1964).

6 Sansonetti, P. J., Kopecko, D. J. & Formal, S. B. Involvement of a plasmid in the invasive ability of Shigella flexneri. Infect Immun 35, 852–860, doi:10.1128/iai.35.3.852-860.1982 (1982).

7 Maurelli, A. T., Baudry, B., d’Hauteville, H., Hale, T. L. & Sansonetti, P. J. Cloning of plasmid DNA sequences involved in invasion of HeLa cells by Shigella flexneri. Infect Immun 49, 164–171 (1985).

8 Carayol, N. & Tran Van Nhieu, G. The inside story of Shigella invasion of intestinal epithelial cells. Cold Spring Harb Perspect Med 3, a016717, doi:10.1101/cshperspect.a016717 (2013).

9 Agaisse, H. Molecular and Cellular Mechanisms of Shigella flexneri Dissemination. Front Cell Infect Microbiol 6, 29, doi:10.3389/fcimb.2016.00029 (2016).

10 Weddle, E. & Agaisse, H. Principles of intracellular bacterial pathogen spread from cell to cell. PLoS Pathog 14, e1007380, doi:10.1371/journal.ppat.1007380 (2018).

11 Sansonetti, P. J., Arondel, J., Fontaine, A., d’Hauteville, H. & Bernardini, M. L. OmpB (osmo-regulation) and icsA (cell-to-cell spread) mutants of Shigella flexneri: vaccine candidates and probes to study the pathogenesis of shigellosis. Vaccine 9, 416–422 (1991).

12 Yum, L. K., Byndloss, M. X., Feldman, S. H. & Agaisse, H. Critical role of bacterial dissemination in an infant rabbit model of bacillary dysentery. Nat Commun 10, 1826, doi:10.1038/s41467-019-09808-4 (2019).

13 Mounier, J., Vasselon, T., Hellio, R., Lesourd, M. & Sansonetti, P. J. Shigella flexneri enters human colonic Caco-2 epithelial cells through the basolateral pole. Infect Immun 60, 237–248, doi:10.1128/iai.60.1.237-248.1992 (1992).

14 Wassef, J. S., Keren, D. F. & Mailloux, J. L. Role of M cells in initial antigen uptake and in ulcer formation in the rabbit intestinal loop model of shigellosis. Infect Immun 57, 858–863, doi:10.1128/iai.57.3.858-863.1989 (1989).

15 Zychlinsky, A., Prevost, M. C. & Sansonetti, P. J. Shigella flexneri induces apoptosis in infected macrophages. Nature 358, 167–169, doi:10.1038/358167a0 (1992).

16 Zychlinsky, A. et al. In vivo apoptosis in Shigella flexneri infections. Infect Immun 64, 5357–5365, doi:10.1128/iai.64.12.5357-5365.1996 (1996).

17 Sansonetti, P. J., Arondel, J., Cavaillon, J. M. & Huerre, M. Role of interleukin-1 in the pathogenesis of experimental shigellosis. J Clin Invest 96, 884–892, doi:10.1172/JCI118135 (1995).

18 Zychlinsky, A., Fitting, C., Cavaillon, J. M. & Sansonetti, P. J. Interleukin 1 is released by murine macrophages during apoptosis induced by Shigella flexneri. J Clin Invest 94, 1328–1332, doi:10.1172/JCI117452 (1994).

19 Perdomo, O. J. et al. Acute inflammation causes epithelial invasion and mucosal destruction in experimental shigellosis. J Exp Med 180, 1307–1319, doi:10.1084/jem.180.4.1307 (1994).

20 Perdomo, J. J., Gounon, P. & Sansonetti, P. J. Polymorphonuclear leukocyte transmigration promotes invasion of colonic epithelial monolayer by Shigella flexneri. J Clin Invest 93, 633–643, doi:10.1172/JCI117015 (1994).

21 Sansonetti, P. J. Microbes and microbial toxins: paradigms for microbial-mucosal interactions III. Shigellosis: from symptoms to molecular pathogenesis. Am J Physiol Gastrointest Liver Physiol 280, G319–323, doi:10.1152/ajpgi.2001.280.3.G319 (2001).

22 Liu, F., Wu, H. Y., Wesselschmidt, R., Kornaga, T. & Link, D. C. Impaired production and increased apoptosis of neutrophils in granulocyte colony-stimulating factor receptor-deficient mice. Immunity 5, 491–501, doi:10.1016/s1074-7613(00)80504-x (1996).

23 Guilbert, L. J. & Stanley, E. R. Specific interaction of murine colony-stimulating factor with mononuclear phagocytic cells. J Cell Biol 85, 153–159, doi:10.1083/jcb.85.1.153 (1980).

24 Bankhead, P. et al. QuPath: Open source software for digital pathology image analysis. Sci Rep 7, 16878, doi:10.1038/s41598-017-17204-5 (2017).

25 Kjeldsen, L., Bainton, D. F., Sengelov, H. & Borregaard, N. Identification of neutrophil gelatinase-associated lipocalin as a novel matrix protein of specific granules in human neutrophils. Blood 83, 799–807 (1994).

26 Elomaa, O. et al. Cloning of a novel bacteria-binding receptor structurally related to scavenger receptors and expressed in a subset of macrophages. Cell 80, 603–609, doi:10.1016/0092-8674(95)90514-6 (1995).

27 Elomaa, O. et al. Structure of the human macrophage MARCO receptor and characterization of its bacteria-binding region. J Biol Chem 273, 4530–4538, doi:10.1074/jbc.273.8.4530 (1998).

28 Kondo, M. et al. Biology of hematopoietic stem cells and progenitors: implications for clinical application. Annu Rev Immunol 21, 759–806, doi:10.1146/annurev.immunol.21.120601.141007 (2003).

29 Mathan, M. M. & Mathan, V. I. Ultrastructural pathology of the rectal mucosa in Shigella dysentery. Am J Pathol 123, 25–38 (1986).

30 Mathan, M. M. & Mathan, V. I. Morphology of rectal mucosa of patients with shigellosis. Rev Infect Dis 13 Suppl 4, S314–318 (1991).

31 Fenton, T. M. et al. Immune Profiling of Human Gut-Associated Lymphoid Tissue Identifies a Role for Isolated Lymphoid Follicles in Priming of Region-Specific Immunity. Immunity 52, 557–570 e556, doi:10.1016/j.immuni.2020.02.001 (2020).

32 Koestler, B. J. et al. Human Intestinal Enteroids as a Model System of Shigella Pathogenesis. Infect Immun 87, doi:10.1128/IAI.00733-18 (2019).

33 Sereny, B. Experimental keratoconjunctivitis shigellosa. Acta Microbiol Acad Sci Hung 4, 367–376 (1957).

34 Voino-Yasenetsky, M. V. & Voino-Yasenetskaya, M. K. Experimental pneumonia caused by bacteria of the Shigella group. Acta Morphol Acad Sci Hung 11, 439–454 (1962).

35 Arm, H. G., Floyd, T. M., Faber, J. E. & Hayes, J. R. Use of Ligated Segments of Rabbit Small Intestine in Experimental Shigellosis. J Bacteriol 89, 803–809, doi:10.1128/jb.89.3.803-809.1965 (1965).

36 Barker, N. et al. Identification of stem cells in small intestine and colon by marker gene Lgr5. Nature 449, 1003–1007, doi:10.1038/nature06196 (2007).

37 Gruber, A. D. et al. Genomic cloning, molecular characterization, and functional analysis of human CLCA1, the first human member of the family of Ca2+-activated Cl-channel proteins. Genomics 54, 200–214, doi:10.1006/geno.1998.5562 (1998).

38 Peeters, T. & Vantrappen, G. The Paneth cell: a source of intestinal lysozyme. Gut 16, 553–558, doi:10.1136/gut.16.7.553 (1975).

39 Talbot, C. & Lytle, C. Segregation of Na/H exchanger-3 and Cl/HCO3 exchanger SLC26A3 (DRA) in rodent cecum and colon. Am J Physiol Gastrointest Liver Physiol 299, G358–367, doi:10.1152/ajpgi.00151.2010 (2010).

40 Metcalfe, C., Kljavin, N. M., Ybarra, R. & de Sauvage, F. J. Lgr5+ stem cells are indispensable for radiation-induced intestinal regeneration. Cell Stem Cell 14, 149–159, doi:10.1016/j.stem.2013.11.008 (2014).

41 van der Post, S. et al. Structural weakening of the colonic mucus barrier is an early event in ulcerative colitis pathogenesis. Gut 68, 2142–2151, doi:10.1136/gutjnl-2018-317571 (2019).

42 Wenneras, C. et al. Blockade of CD14 increases Shigella-mediated invasion and tissue destruction. J Immunol 164, 3214–3221, doi:10.4049/jimmunol.164.6.3214 (2000).

43 Raqib, R. et al. Innate immune responses in children and adults with Shigellosis. Infect Immun 68, 3620–3629, doi:10.1128/IAI.68.6.3620-3629.2000 (2000).

44 Hackert, N. S. et al. Human and mouse neutrophils share core transcriptional programs in both homeostatic and inflamed contexts. Nat Commun 14, 8133, doi:10.1038/s41467-023-43573-9 (2023).

45 Gautier, E. L. et al. Gene-expression profiles and transcriptional regulatory pathways that underlie the identity and diversity of mouse tissue macrophages. Nat Immunol 13, 1118–1128, doi:10.1038/ni.2419 (2012).

46 Ley, K., Laudanna, C., Cybulsky, M. I. & Nourshargh, S. Getting to the site of inflammation: the leukocyte adhesion cascade updated. Nat Rev Immunol 7, 678–689, doi:10.1038/nri2156 (2007).

47 Raqib, R. et al. Persistence of local cytokine production in shigellosis in acute and convalescent stages. Infect Immun 63, 289–296, doi:10.1128/iai.63.1.289-296.1995 (1995).

48 Kolaczkowska, E. & Kubes, P. Neutrophil recruitment and function in health and inflammation. Nat Rev Immunol 13, 159–175, doi:10.1038/nri3399 (2013).

49 Flannagan, R. S., Jaumouille, V. & Grinstein, S. The cell biology of phagocytosis. Annu Rev Pathol 7, 61–98, doi:10.1146/annurev-pathol-011811-132445 (2012).

50 Eislmayr, K. D. et al. Macrophages orchestrate elimination of Shigella from the intestinal epithelial cell niche via TLR-induced IL-12 and IFN-gamma. Cell Host Microbe 33, 1535–1549 e1537, doi:10.1016/j.chom.2025.08.001 (2025).

51 Wang, F. et al. RNAscope: a novel in situ RNA analysis platform for formalin-fixed, paraffin-embedded tissues. J Mol Diagn 14, 22–29, doi:10.1016/j.jmoldx.2011.08.002 (2012).

52 Hall, C. P., Jadeja, N. B., Sebeck, N. & Agaisse, H. Characterization of MxiE– and H-NS-Dependent Expression of ipaH7.8, ospC1, yccE, and yfdF in Shigella flexneri. mSphere 7, e0048522, doi:10.1128/msphere.00485-22 (2022).

53 Hubbard, T. et al. The Ensembl genome database project. Nucleic Acids Res 30, 38–41, doi:10.1093/nar/30.1.38 (2002).

54 Love, M. I., Huber, W. & Anders, S. Moderated estimation of fold change and dispersion for RNA-seq data with DESeq2. Genome Biol 15, 550, doi:10.1186/s13059-014-0550-8 (2014).

55 Subramanian, A. et al. Gene set enrichment analysis: a knowledge-based approach for interpreting genome-wide expression profiles. Proc Natl Acad Sci U S A 102, 15545–15550, doi:10.1073/pnas.0506580102 (2005).

